# ISRIB promotes aggregation of TIA-1 by modulating its phase separation *in vitro*

**DOI:** 10.1101/2025.07.07.663586

**Authors:** Miyu Murata, Hitomi Kimura, Shin-ichi Tate, Kyota Yasuda

## Abstract

Stress granules (SGs) are non-membranous biomolecular condensates formed through liquid-liquid phase separation (LLPS) of RNA-binding proteins (RBPs) and RNA under stress conditions. T-cell intracellular antigen-1 (TIA-1), a major RBP of SGs, can undergo LLPS via its low-complexity domain, contributing to SG nucleation. Integrated stress response inhibitor (ISRIB), a small molecule that enhances eIF2B activity and inhibits the integrated stress response, has been widely studied for its therapeutic potential in neurodegenerative diseases. However, little is known about how ISRIB directly affects the behavior of SG proteins. Here, we show that ISRIB enhances TIA-1 phase separation and promotes its aggregation in vitro. Interestingly, this effect was mitigated in the presence of RNA or cell lysates, suggesting that RNA-binding plays a protective role against ISRIB-induced aggregation. These findings imply that ISRIB alters the physical properties of SG in an RNA-dependent manner, raising important considerations for their therapeutic application.

**Graphic abstract:** 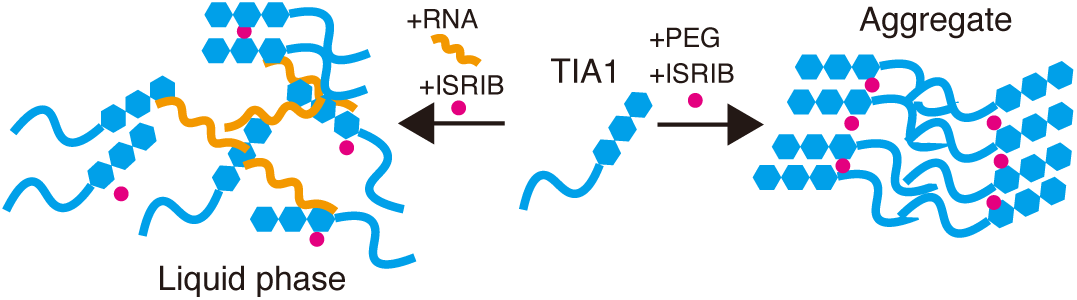

## 1. Introduction

Biomolecular condensates, including stress granules (SGs), are membrane-less organelles formed via liquid-liquid phase separation (LLPS), a process driven by multivalent, weak interactions between intrinsically disordered regions of proteins and RNA (1, 2). SGs assemble dynamically in response to various stress stimuli and sequester untranslated mRNAs and RNA-binding proteins (RBPs), thereby modulating translation, preserving cellular homeostasis, and facilitating stress adaptation (3).

Among the SG-associated RBPs, T-cell intracellular antigen-1 (TIA-1) plays a central role in SG nucleation (4, 5). TIA-1 contains three RNA recognition motifs (RRMs) and a low-complexity domain (LCD), both of which contribute to its ability to undergo LLPS and form dynamic granules (4, 6). Furthermore, LCD-mediated interactions are especially sensitive to physicochemical conditions and post-translational modifications, which makes TIA-1 highly responsive to environmental cues (7, 8). However, under chronic or excessive stress, this response may lead to the aggregation of pathological proteins. Aberrant phase transitions of RBPs, such as TIA-1, have been implicated in the pathogenesis of neurodegenerative diseases, such as amyotrophic lateral sclerosis (ALS) and frontotemporal dementia (FTD) (9–11).

Modulation of LLPS by small molecules has emerged as a promising approach to influence SG dynamics and potentially correct the pathological aggregation behavior. Integrated stress response inhibitor (ISRIB) is one such molecule that has garnered significant attention. ISRIB enhances the activity of eIF2B, a guanine nucleotide exchange factor essential for initiating protein translation (12). During cellular stress, phosphorylation of eIF2α inhibits eIF2B, suppressing translation and promoting SG formation (13). ISRIB counteracts this by reactivating eIF2B even in the presence of phosphorylated eIF2α, thereby restoring translation and suppressing SG assembly (14). This mechanism makes ISRIB a promising therapeutic candidate for various neurological conditions including traumatic brain injury and ALS (15).

Although intracellular mechanisms of ISRIB are well-documented, their direct interactions with SG components remain underexplored. Interestingly, prior research has shown that GFP-G3BP1 droplets isolated from ISRIB-treated cells exhibit an increased size, raising the possibility that ISRIB might modulate the physical properties of SG proteins directly, independent of integrated stress response (ISR) suppression (16). Understanding whether ISRIB exerts such direct effects on individual SG components is crucial to determine its broader biological impact and therapeutic safety.

In this study, we investigated the effects of ISRIB on TIA-1 phase behavior in both cellular and in vitro systems. We found that while ISRIB universally promotes TIA-1 condensation, its ability to induce aggregation is context-dependent, occurring only under RNA-deficient conditions. These findings highlight the importance of RNA in safeguarding against aberrant phase transitions and suggest that ISRIB, despite its therapeutic benefits, may pose aggregation risks in environments with compromised RNA availability.

## 2. Materials and Methods

### 2.1. Cell Culture and Stress Induction

HEK293T cells (cat# RCB2202; RIKEN Cell Bank, Japan; RRID: CVCL_0063) were cultured in high-glucose Dulbecco’s Modified Eagle Medium (DMEM; cat# C11995500CP, Gibco, USA) supplemented with 10% fetal bovine serum, 100 U/mL penicillin, and 100 µg/mL streptomycin (cat# 15140122, Gibco). Cells were treated with 500 µM sodium arsenite (cat# S7400, Sigma-Aldrich, USA) for 1 h to induce stress granule formation. For the ISRIB treatment, ISRIB was added to the culture medium at a final concentration of 200 nM, and control samples were treated with an equivalent volume of DMSO. For the Cycloheximide (CHX) treatment, CHX was added to the culture medium at a final concentration of 100 μg/ml.

### 2.2. Plasmid Construction and Transfection

The TIA-1 coding sequence (full-length or RNA recognition motifs, RRMs: 1–290 aa) was cloned into the pET28a/mNeonGreen (mNG) vector using BamHI and EcoRI restriction sites. For mammalian expression, GFP-TIA-1 was constructed by subcloning EGFP-C1 using SacI and BamHI. Transient transfection into HEK293T cells was performed using PEI Max (cat# 24765-100, Polysciences, Inc., USA) according to the manufacturer’s instructions. Cells were used at 36–48 h post-transfection.

### 2.3. Isolation of the SG-Enriched Fraction

SG-enriched fractions were isolated as previously described with modifications (17). HEK293T cells cultured in 10-cm dishes were treated with 500 µM arsenite for 1 h, washed with PBS, and collected into 6 mL of lysis buffer (50 mM Tris-HCl pH 7.4, 100 mM potassium acetate, 2 mM magnesium acetate, 0.5 mM DTT, 50 µg/mL heparin, 0.5% NP-40, protease inhibitor cocktail). Cells were lysed on ice for 10 min, and the lysate was centrifuged at 1,000 × *g* for 5 min at 4 °C to remove debris. The supernatant was further centrifuged at 18,000 × *g* for 20 min at 4 °C. The resulting pellet was resuspended in 600 µL lysis buffer and centrifuged at 900 × *g* for 2 min at 4 °C. The final supernatant was used as an SG-rich fraction. ISRIB (200 nM final concentration) or DMSO was added, and the samples were immediately used for further experiments.

### 2.4. Immunofluorescence Staining

Cells were cultured on fibronectin-coated cover glasses (cat# 0111580; Marienfeld, Germany) and fixed in 4% paraformaldehyde in PBS for 15 min. Cells were permeabilized with 0.2% Triton X-100 in PBS for 5 min, blocked with 5% fetal bovine serum in PBS for 1 h at room temperature, and incubated overnight at 4 °C with primary antibody anti-G3BP1 (cat# ab56574, Abcam, UK). After washing, an Alexa Fluor 647-conjugated secondary antibody (cat# A-21244, Invitrogen, Carlsbad, CA, USA) was added for 1 h at room temperature. Coverslips were mounted using Fluoromount-G containing DAPI (cat# D11130H, Matsunami, Japan).

### 2.5. Turbidity Assay

Turbidity was measured using a TriStar microplate reader (Berthold Technologies). The SG fractions or purified protein samples (with or without ISRIB) were dispensed into 96-well plates in triplicate. The optical density was measured at 525 nm (OD525). The average absorbance from triplicate wells was used as the experimental data.

### 2.6. Imaging and Quantification

Images were acquired using a spinning disk confocal microscope (IX83, Olympus) equipped with a CSU-W1 scanner (Yokogawa) and a 100× oil objective lens (NA 1.4, UPLSAPO 100XO) or a 20× lens (NA 0.75, UPLSAPO 20X). GFP and mNeonGreen (mNG) were excited at 488 nm, and Alexa Fluor 647 was excited at 637 nm. Z-stacks were acquired (0.2 µm steps × 40 slices), and projection images were generated using Fiji (ImageJ) (18). Granules were quantified by setting fluorescence intensity thresholds, and granule area and circularity were calculated based on calibrated pixel dimensions.

### 2.7. Recombinant Protein Expression and Purification

The pET28a plasmid encoding mNG-TIA-1 (full-length or RRM) was transfected into *Escherichia coli* BL21(DE3) cells (cat# DS250, BioDynamics Laboratory Inc.). Cultures were grown in LB medium with 20 µg/mL kanamycin at 37 °C to OD600 ≈ 0.6, then induced with 1 mM IPTG and incubated at 16 °C for 16–20 h. The cells were harvested, resuspended in buffer A (50 mM Tris-HCl, pH 9.0, 200 mM NaCl, 2 M urea, and 20 mM imidazole), lysed by sonication, and clarified by centrifugation. The supernatant was loaded onto a Ni-NTA column (GE Healthcare), washed with buffers A, B (4 M NaCl), and C (50 mM NaCl), and eluted with buffer E (500 mM imidazole). Proteins were dialyzed against HEPES (20 mM, pH 7.4) and Tris-HCl (50 mM, pH 7.5, and 500 mM arginine) and quantified using a NanoDrop spectrophotometer.

### 2.8. In Vitro Droplet Formation Assays

The recombinant mNG-TIA-1 protein was adjusted to be 5 μM in HEPES buffer (20 mM, pH 7.4), incubated for 30 min, and subjected to LLPS induction by the addition of 10% PEG-3350 (cat# 202444PEG-3350; Sigma-Aldrich) or total RNA (50 ng/μl at final concentration) extracted from HEK293T cells. ISRIB or DMSO was added and incubated for 30 min before observation. ZnCl^2^ (3 μM at final concentration) was used as another inducer of phase separation.

### 2.9. Statistical Analysis

Data were analyzed using Prism 10 (GraphPad Software). Shapiro–Wilk tests were used to assess normality. Non-parametric or parametric tests, including unpaired t-tests and Dunnett’s multiple comparison tests, were applied accordingly. Statistical significance was defined as *P* < 0.05.

## 3. Results

### 3.1. ISRIB promotes SG clearance in cells

To validate the reported cellular effects of ISRIB, we first confirmed that ISRIB inhibited SG formation in HEK293T cells. Upon sodium arsenite treatment, SGs were robustly formed, as visualized by G3BP1 immunostaining. ISRIB treatment reduced SG formation and promoted SG disassembly (Supplementary Fig. 1A–D), consistent with previous reports.

To test whether ISRIB acts on pre-existing SGs, cells were first exposed to arsenite for 1 hour to induce SG formation, followed by ISRIB treatment. Microscopy and quantification confirmed that ISRIB effectively promoted the clearance of SGs even after their initial assembly (Supplementary Fig. 1E-H), supporting a broader temporal window for ISRIB activity.

To evaluate whether the SG clearance effect of ISRIB involves translation-dependent mechanisms, we also examined the effect of cycloheximide (CHX), a classical translation inhibitor, on arsenite-induced SGs. CHX treatment after arsenite-induced SG formation reduced both SG number and size to a similar extent as ISRIB treatment. Interestingly, co-treatment with CHX and ISRIB resulted in an even greater reduction in SG number and size compared to ISRIB alone (Supplementary Fig. 2), suggesting partial additive or synergistic effects. These findings indicate that ISRIB and CHX may promote SG clearance through both overlapping and distinct mechanisms.

### 3.2. ISRIB promotes condensation of TIA-1 in cell lysate

Next, we examined the effect of ISRIB on the SG-enriched fraction isolated from the arsenite-treated cells expressing GFP-TIA-1 (Fig. 1A). The isolated SG fraction displayed droplet-like GFP-TIA-1 condensates under the microscope (Fig. 1B). Notably, ISRIB treatment led to increased turbidity (Fig. 1C), larger droplet size (Fig. 1D), and higher granule count (Fig. 1E) compared to the DMSO-treated controls, consistent with previous *ex vivo* observations using GFP-G3BP1 (16).

**Figure 1.**
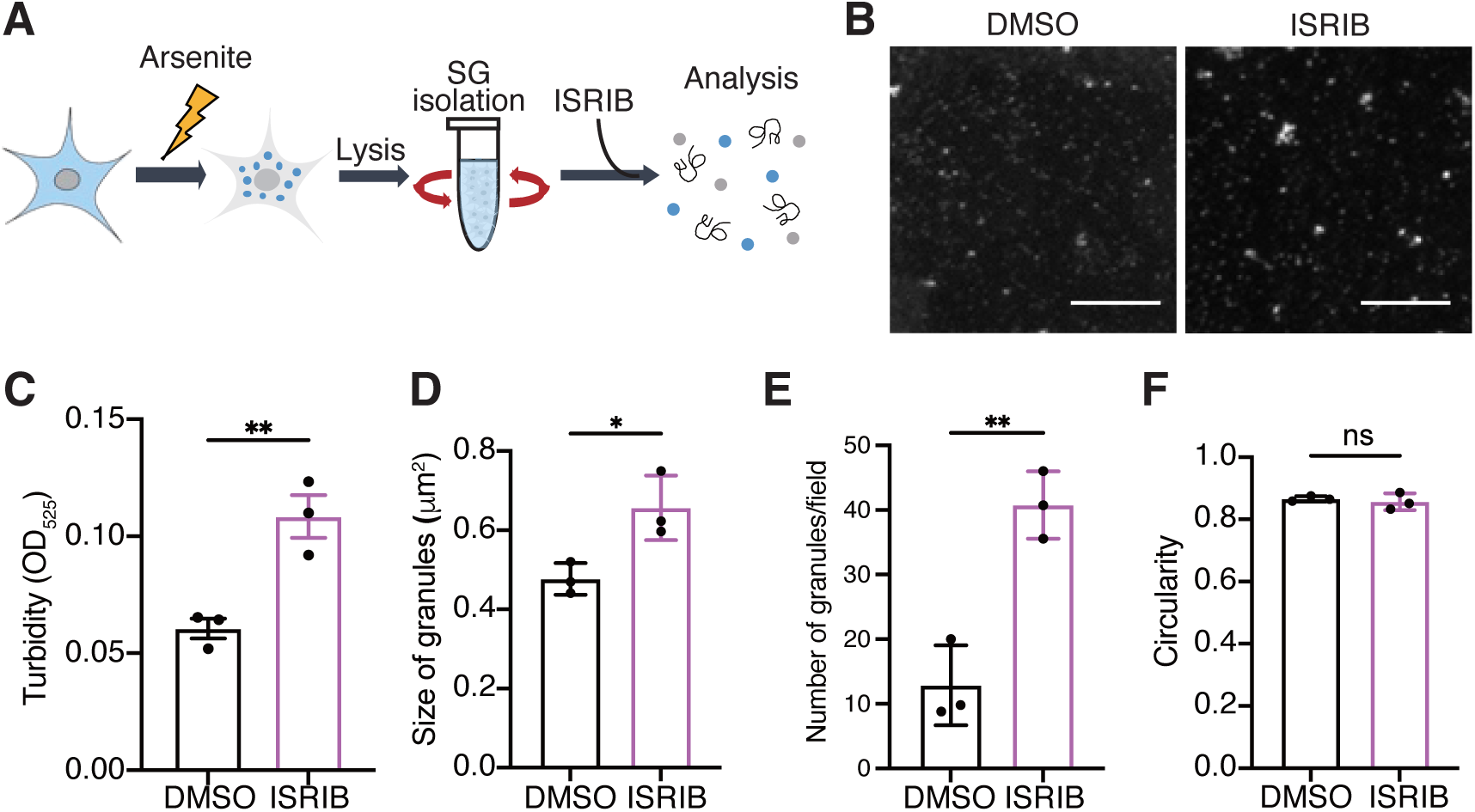
ISRIB induces the LLPS of TIA-1 in the isolated SG core fraction. **(A)** Schematic view of the experiment. HEK293T cells were exposed to arsenite stress for 1 h to induce the formation of SGs. The cells were then lysed and centrifuged to isolate the SG fraction. ISRIB was added to the SG fraction, and samples were either observed under a confocal microscope or subjected to turbidity measurements. **(B)** Representative images of GFP-TIA-1 in the SG fraction of cell lysate with and without ISRIB treatment. Control samples were treated with DMSO, the vehicle of ISRIB. Bars indicate 10 μm. **(C)** Turbidity of the indicated samples was plotted as a bar graph. Each data point indicates the average of triplicated samples. **(D)** The size of granules analyzed from images of the indicated samples was plotted as a bar graph. Each data point represents the average in one trial. *n(granule)* = 222 (DMSO) and 810 (ISRIB) in total. **(E)** The number of granules in an average number of granules per field analyzed from images of the indicated samples was plotted as a bar graph. Each data point represents the average in one trial. *n(field)* = 17 (DMSO) and 20 (ISRIB) in total. **(F)** The circularity of granules analyzed from images of the indicated samples was plotted as a bar graph. *n(granule)* = 222 (DMSO) and 810 (ISRIB) in total. All data points were collected from independent trials. Results of the control (DMSO) and experimental (ISRIB) samples were compared by using an unpaired t-test. \*\**P* < 0.01 and \**P* < 0.05.

However, the circularity remained largely unaffected (Fig. 1F). Because circularity serves as an indicator of droplet morphology and internal dynamics—with lower circularity implying deformation and potential aggregation—this result suggests that ISRIB enhanced TIA-1 condensation without promoting aggregation under these lysate-containing conditions.

### 3.3. ISRIB promotes the aggregation of recombinant TIA-1

To assess whether ISRIB directly promoted TIA-1 phase separation, we used recombinant mNG-TIA-1 protein purified from *E. coli* and induced LLPS in vitro using 10% PEG-3350 as a molecular crowding agent. Turbidity measurements revealed a concentration-dependent increase upon ISRIB treatment (Fig. 2A), suggesting that ISRIB facilitated the formation of condensed protein assemblies.

**Figure 2.**
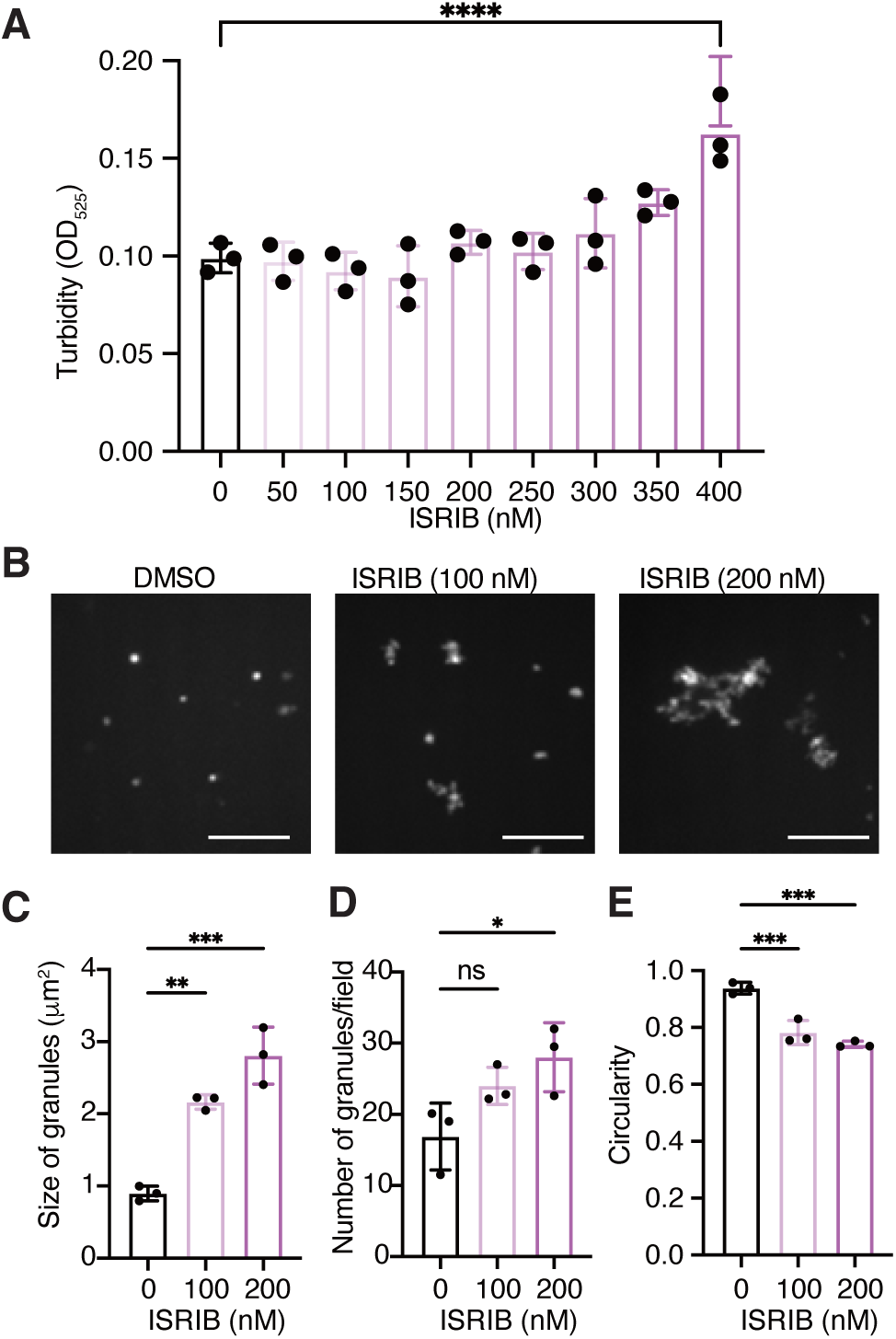
ISRIB promotes the aggregation of PEG-induced LLPS of synthesized TIA-1. **(A)** The turbidity of the synthesized mNG-TIA-1 phase separated with PEG with indicated ISRIB concentrations was plotted as a bar graph. Each data point indicates the average of triplicated samples. **(B)** Representative images of synthesized mNG-TIA-1 phase separated with PEG with and without the indicated concentration of ISRIB. Control samples were treated with DMSO. Bars indicate 10 μm. **(C)** The size of granules analyzed from images of the indicated samples was plotted as a bar graph. Each data point represents the average in one trial. *n(granule)* = 344 (DMSO), 486 (100 nM, ISRIB), and 550 (200 nM, ISRIB) in total. **(D)** The number of granules in an average number of granules per field analyzed from images of the indicated samples was plotted as a bar graph. Each data point represents the average in one trial. *n(field)* = 20 (DMSO), 20 (100 nM, ISRIB), and 20 (200 nM, ISRIB) in total. **(E)** The circularity of granules analyzed from images of the indicated samples was plotted as a bar graph. *n(granule)* = 344 (DMSO), 486 (100 nM, ISRIB), and 550 (200 nM, ISRIB) in total. All data points were collected from independent trials. Results of control (DMSO) and experimental (ISRIB) samples were compared by using Dunnett’s multiple comparisons test. \**P* < 0.05, \*\**P* < 0.005, and \*\*\**P* < 0.0005.

Fluorescence microscopy corroborated these findings; ISRIB-treated samples exhibited markedly larger and more numerous droplets than the DMSO controls (Fig. 2B–D). Furthermore, droplet circularity decreased significantly with increasing ISRIB concentration (Fig. 2E), indicating a transition from dynamic, spherical condensates to more irregular, solid-like aggregates. These observations collectively support a direct role of ISRIB in promoting TIA-1 condensation and aggregation, independent of its canonical ISR-related function.

To assess whether the aggregation-promoting effect of ISRIB was specific to PEG-induced LLPS or generalizable to other LLPS inducers, we performed similar assays using Zn^2+^ as an alternative induction agent (19, 20). In Zn^2+^-induced LLPS conditions, ISRIB treatment increased droplet size and decreased circularity, consistent with PEG results (Sup Fig. 3). These findings indicate that ISRIB promotes aggregation-prone transitions across distinct LLPS induction mechanisms.

It is noteworthy that the concentration range in which ISRIB increased turbidity was higher than that required to alter granule morphology. This discrepancy likely reflects differences in assay sensitivity: turbidity measurements capture total condensed mass, while microscopy-based parameters—such as droplet size, number, and circularity—can detect earlier or more subtle physical transitions. These results underscore the importance of using multiple complementary readouts when evaluating phase separation dynamics.

### 3.4. RNA inhibits ISRIB-induced aggregation of TIA-1

The contrasting effects observed in Fig. 1 and Fig. 2 suggested that cellular factors may suppress ISRIB-induced aggregation. In Fig. 1, using SG-enriched lysates containing endogenous RNA, ISRIB enhanced TIA-1 condensation without altering droplet circularity, implying that aggregation was prevented. In contrast, ISRIB treatment of recombinant TIA-1 under PEG-induced LLPS conditions promoted aggregation-like transitions, as evidenced by decreased circularity. Given that RNA is a major SG component, we hypothesized that it might play a protective role against ISRIB-induced aggregation.

To test this hypothesis, we compared PEG-induced LLPS with RNA-induced LLPS using total RNA extracted from HEK293T cells (Fig. 3A, B). ISRIB markedly increased granule size in PEG-induced LLPS but had no significant effect under RNA-induced conditions (Fig. 3C). Granule number remained unchanged in both conditions (Fig. 3D). Importantly, ISRIB significantly decreased circularity in PEG-induced LLPS but not in RNA-induced LLPS (Fig. 3E), indicating that RNA stabilizes TIA-1 condensates in a liquid-like state and suppresses ISRIB-induced aggregation.

**Figure 3.**
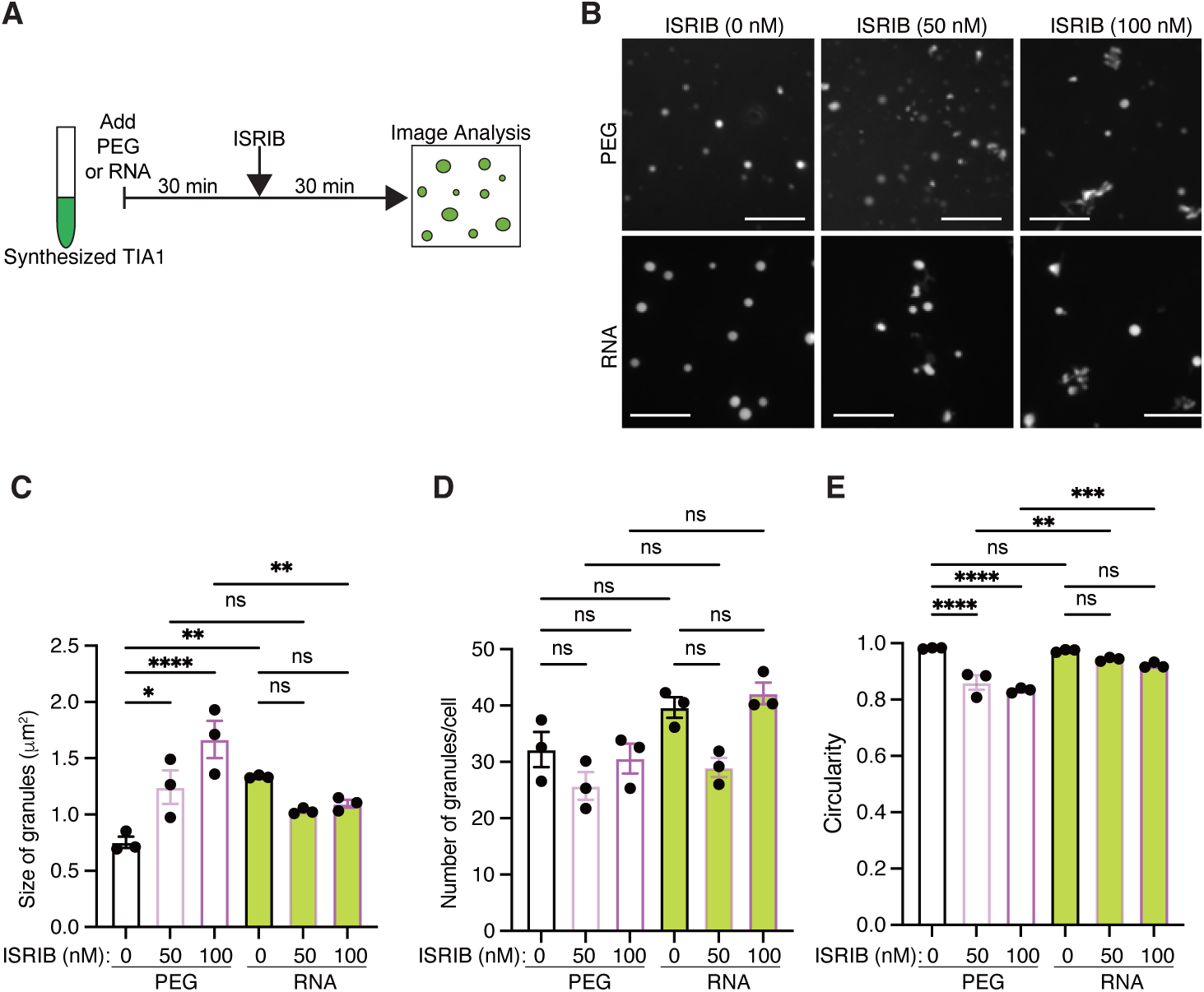
Pre-RNA treatment prevents ISRIB-driven aggregation of TIA-1 *in vitro*. **(A)** Schematic view of the experiment. *E. coli-synthesized* and purified mNG-TIA-1 was phase-separated with PEG or RNA. ISRIB was added 30 minutes later, and samples were observed under a confocal microscope after an additional 30 minutes. **(B)** Representative images of synthesized mNG-TIA-1 treated as in (A). Bars indicate 10 μm. **(C)** The size of granules analyzed from images of synthesized mNG-TIA-1, phase separated with PEG (white body) or RNA (green body), was plotted as a bar graph. Each data point represents the average in one trial. *n(granule)* = 676 (PEG, DMSO), 591 (PEG, 50 nM ISRIB), 691 (PEG, 100 nM ISRIB), 873 (RNA, DMSO), 680 (RNA, 50 nM ISRIB), and 839 (RNA, 100 nM ISRIB) in total. **(D)** The number of granules analyzed from images of synthesized mNG-TIA-1, phase separated with PEG (white body) or RNA (green body), was plotted as a bar graph. Each data point represents the average in one trial. *n(field)* = 21 (PEG, DMSO), 23 (PEG, 50 nM ISRIB), 23 (PEG, 100 nM ISRIB), 22 (RNA, DMSO), 20 (RNA, 50 nM ISRIB), and 22 (RNA, 100 nM ISRIB) in total. **(E)** The circularity of granules analyzed from images of synthesized mNG-TIA-1, phase separated with PEG (white body) or RNA (green body), was plotted as a bar graph. Each data point represents the average in one trial. *n(granule)* = 676 (PEG, DMSO), 591 (PEG, 50 nM ISRIB), 691 (PEG, 100 nM ISRIB), 873 (RNA, DMSO), 680 (RNA, 50 nM ISRIB), and 839 (RNA, 100 nM ISRIB) in total. All data points were collected from independent trials. Results of control (DMSO) and experimental (ISRIB) samples were compared by using Dunnett’s multiple comparisons test. \**P* < 0.05, \*\**P* < 0.005, \*\*\**P* < 0.0005, and \*\*\*\**P* < 0.0001.

To further assess whether RNA could reverse ISRIB-induced aggregation, we added RNA after ISRIB treatment in PEG-induced LLPS assays (Fig. 4A). Interestingly, RNA addition after ISRIB failed to restore droplet circularity or reduce granule size (Fig. 4C– E). Instead, circularity decreased further, suggesting that once aggregation-prone transitions are established by ISRIB, subsequent RNA addition cannot revert these structural changes.

**Figure 4.**
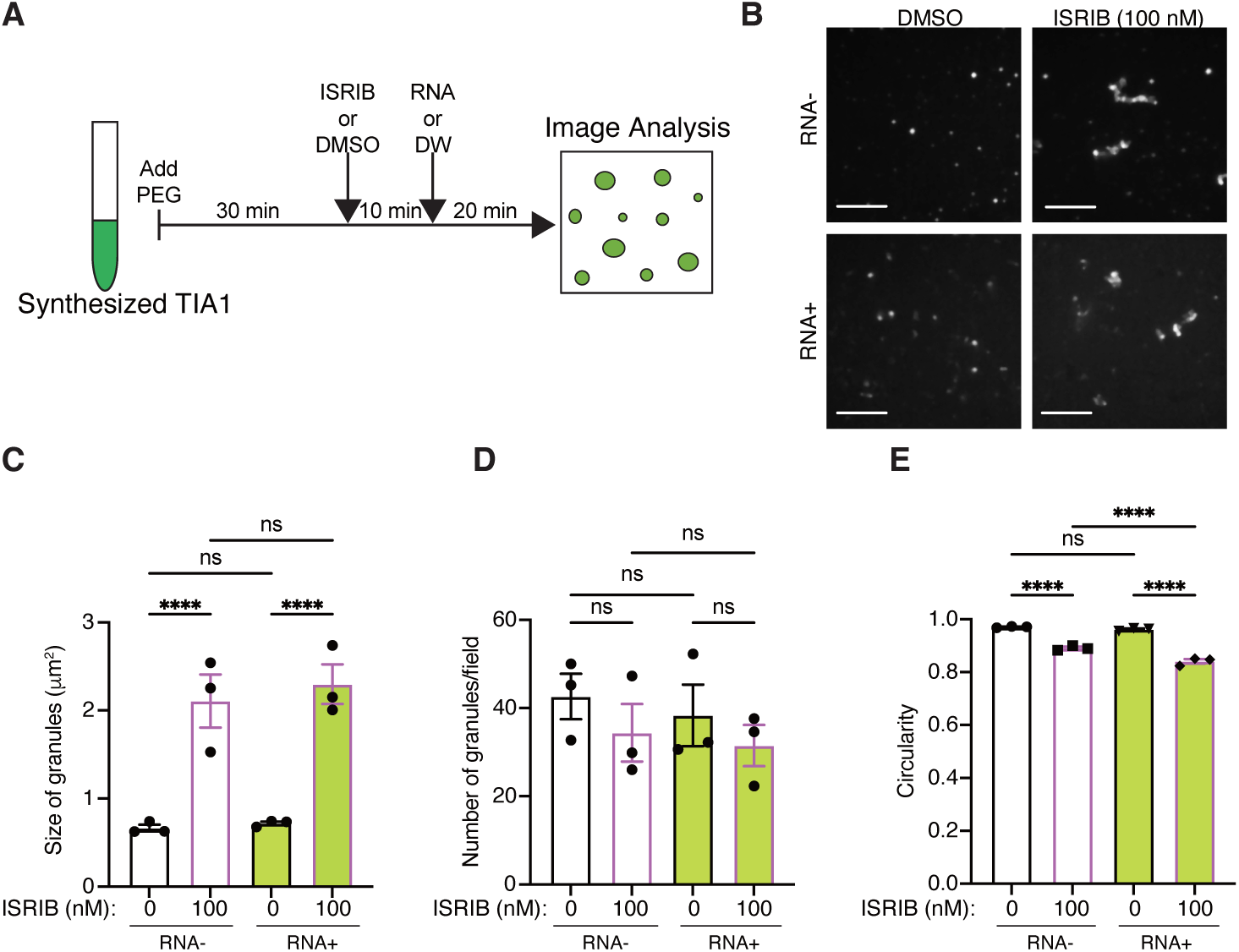
ISRIB-driven aggregation of TIA-1 cannot be recovered *in vitro*. **(A)** Schematic view of the experiment. *E. coli-synthesized* and purified mNG-TIA-1 was phase-separated with PEG. ISRIB was added 30 minutes later, and RNAs were added 10 minutes later. Then samples were observed under a confocal microscope after an additional 20 minutes. **(B)** Representative images of observed mNG-TIA-1 samples treated as in (A). Control samples were treated with DMSO. Bars indicate 10 μm. **(C)** The size of granules analyzed from images of synthesized mNG-TIA-1, treated with (green body) or without RNA (white body), was plotted as a bar graph. Each data point represents the average in one trial. *n(granule)* = 2859 (RNA-, DMSO), 1932 (RNA-, 100 nM ISRIB), 851 (RNA+, DMSO), and 745 (RNA+, 100 nM ISRIB) in total. **(D)** The number of granules analyzed from images of synthesized mNG-TIA-1, treated with (green body) or without RNA (white body), was plotted as a bar graph. Each data point represents the average in one trial. *n(granule)* = 30 (RNA-, DMSO), 30 (RNA-, 100 nM ISRIB), 30 (RNA+, DMSO), and 30 (RNA+, 100 nM ISRIB) in total. **(E)** The circularity of granules analyzed from images of synthesized mNG-TIA-1, treated with (green body) or without RNA (white body), was plotted as a bar graph. Each data point represents the average in one trial. *n(granule)* = 2859 (RNA-, DMSO), 1932 (RNA-, 100 nM ISRIB), 851 (RNA+, DMSO), and 745 (RNA+, 100 nM ISRIB) in total. All data points were collected from independent trials. Results were compared by using Dunnett’s multiple comparisons test. \*\*\*\**P* < 0.0001.

### 3.5. ISRIB promotes aggregation under Zn²⁺-induced LLPS, and RNA post-addition fails to reverse aggregation

Finally, to determine whether ISRIB’s aggregation-promoting effect is specific to PEG-induced LLPS or generalizable to other LLPS mechanisms, we performed similar assays using Zn²⁺ as an alternative LLPS inducer (Sup Fig. 3). ISRIB treatment under Zn²⁺-induced conditions increased droplet size and decreased circularity, mirroring results obtained with PEG-induced LLPS (Sup Fig. 3B-D).

Furthermore, we tested whether RNA could reverse ISRIB-induced aggregation when added after granule formation. Interestingly, RNA addition after ISRIB treatment did not suppress aggregation in Zn²⁺-induced droplets (Sup Fig. 3D). Instead, circularity decreased further, indicating enhanced irregularity and reduced fluidity. This suggests that while RNA can protect against ISRIB-induced aggregation when present during LLPS initiation (as seen in PEG assays), it is insufficient to reverse aggregation-prone states once established.

Together, these findings demonstrate that ISRIB promotes TIA-1 aggregation across distinct LLPS induction mechanisms, and that the modulatory effects of RNA are highly context- and timing-dependent.

## 4. Discussion

### 4.1. ISRIB exhibits dual roles in TIA-1 phase behavior

ISRIB is classically known as a translational regulator that suppresses SG formation by enhancing eIF2B activity, thereby restoring translation suppressed by phosphorylated eIF2α (12, 13, 21, 22). Consistent with previous reports, we confirmed that ISRIB not only inhibits SG formation but also promotes SG clearance when administered after SG assembly in HEK293T cells (Sup Fig. 1). Given its ability to clear stress granules, ISRIB has been considered a promising therapeutic agent for neurodegenerative diseases such as ALS, where persistent and aberrant SGs contribute to pathological protein aggregation (6, 23).

Interestingly, our assays revealed an additional and unexpected role of ISRIB: its effect on TIA-1 aggregation was highly context-dependent. In cell lysate experiments (Fig. 1), which contain abundant RNA, ISRIB promoted condensation as indicated by increased turbidity, droplet size, and number, but did not reduce circularity, suggesting no aggregation under these RNA-rich conditions. In contrast, under PEG-induced RNA-deficient conditions (Fig. 2), ISRIB treatment led to decreased circularity, indicating aggregation promotion. To test whether this aggregation-promoting effect was specific to PEG-induced LLPS, we employed Zn²⁺ as an alternative LLPS inducer. Similar to PEG, ISRIB treatment under Zn²⁺-induced conditions increased granule size and decreased circularity (Sup Fig. 3). Notably, previous studies have shown that Zn²⁺ binds to the RRM2 domain of TIA-1, promoting its multimerization and condensation (20), implying that ISRIB promotes aggregation across distinct LLPS induction mechanisms.

Furthermore, in RNA-induced LLPS (Fig. 3), ISRIB failed to induce aggregation-like transitions, highlighting RNA’s protective role in stabilizing TIA-1 condensates in a liquid-like state. However, this protective effect was timing-dependent; RNA addition after ISRIB-induced aggregation did not reverse droplet solidification and even exacerbated irregular morphology (Fig. 4, Sup Fig. 3). This indicates that once TIA-1 aggregation has been promoted by ISRIB, subsequent RNA binding is insufficient to remodel aggregates back to dynamic states.

In our analyses, changes in SG number, size, and circularity provide insights into the physiological state of condensates. An increase in SG number suggests enhanced nucleation, while an increase in size indicates droplet growth via fusion or component recruitment. Reduced circularity indicates decreased internal fluidity, suggesting a transition toward solid-like, aggregation-prone states, which is often associated with pathological protein aggregation. In this context, ISRIB promoted increases in droplet size and number with decreased circularity under RNA-deficient in vitro conditions, indicating promotion of aggregation. By contrast, in cells, ISRIB reduced SG size and number, suggesting suppression or clearance of SGs. Thus, these parameters collectively highlight the context-dependent dual effects of ISRIB on TIA-1 phase behavior.

Collectively, these results demonstrate ISRIB’s context-dependent duality: it acts as a protective translational modulator in cells with intact RNA homeostasis but may promote pathogenic aggregation under RNA-deficient or stress-altered conditions. Such scenarios are conceivable in diseases like ALS or FTD, where RNA buffering capacity is thought to be compromised due to altered RNA-binding protein function (6, 24).

### 4.2. Mechanistic implications of ISRIB-induced aggregation

The precise mechanism by which ISRIB promotes TIA-1 aggregation remains unclear. One possibility is direct binding to TIA-1 RRM domains, competing with RNA and enhancing protein–protein interactions. Alternatively, ISRIB may indirectly modulate the physicochemical properties of the environment—such as dielectric constant or hydrophobicity—to favor LLPS and aggregation. Similar indirect modulation has been reported for small molecules like ATP, which regulate LLPS without direct target binding (25).

Given that ISRIB promoted aggregation under both PEG- and Zn²⁺-induced LLPS conditions, its action likely generalizes across different biophysical contexts. This suggests that ISRIB could exacerbate the aggregation tendencies of RBPs under pathological conditions involving metal dyshomeostasis or RNA metabolism impairment.

Further structural studies will be necessary to identify potential ISRIB binding sites on SG proteins, and physicochemical assays may clarify whether ISRIB alters condensate properties via indirect environmental effects.

### 4.3. Physiological relevance and context-dependent duality

In our in vitro assays, PEG was used as a macromolecular crowding agent to mimic the crowded intracellular environment, facilitating LLPS by volume exclusion effects, although PEG itself is not a physiological component. In cells, macromolecular crowding arising from high concentrations of proteins, nucleic acids, and complexes creates an environment where phase separation is thermodynamically favored. Although PEG itself is an artificial agent, it replicates the excluded volume effects and crowded conditions found in the cytoplasm. Therefore, PEG-induced LLPS is considered a biophysically relevant in vitro model to study phase behavior under cellular-like crowded environments, while RNA-induced LLPS models the specific molecular interactions that occur in RNA-rich contexts such as stress granules.

Our cell-based assays suggest that ISRIB’s SG clearance effect may not be solely attributed to its role in restoring translation. While cycloheximide (CHX), a translation inhibitor, also reduced SG number and size, co-treatment with ISRIB and CHX resulted in an even greater reduction, suggesting partial additive or synergistic effects. This implies that ISRIB may promote SG clearance through both translation-dependent mechanisms and translation-independent pathways, possibly by directly modulating the phase separation dynamics of SG components such as TIA-1.

These seemingly contradictory effects of ISRIB—promoting TIA-1 aggregation in vitro under RNA-deficient conditions, while inhibiting SG formation or promoting SG disassembly in RNA-rich cellular contexts—can be reconciled by considering its dual mechanisms of action. In cells, ISRIB enhances eIF2B activity to restore translation, thereby reducing the need for SG assembly. Furthermore, the abundant RNA present in cells acts as a molecular buffer, stabilizing TIA-1 in a liquid-like condensate state and preventing ISRIB-induced aggregation. By contrast, in vitro experiments lack both translation machinery and sufficient RNA buffering capacity, thereby unmasking ISRIB’s direct physicochemical effect on TIA-1, which promotes phase transition into more aggregation-prone states. Thus, the overall outcome of ISRIB treatment depends on the cellular context, particularly RNA availability and the presence of translational regulation pathways, highlighting its context-dependent duality with potential pathological implications.

### 4.4. Therapeutic considerations and future directions

While ISRIB remains a promising therapeutic candidate due to its neuroprotective effects in animal models (15, 26), our findings raise caution regarding its potential to promote protein aggregation under disease-altered cellular conditions. Future studies should:

- Identify ISRIB-protein binding interfaces using structural and biochemical approaches;
- Evaluate ISRIB effects in disease models with altered RNA homeostasis or SG dynamics to define its therapeutic window; and
- Explore combinatorial strategies, such as co-administration with RNA mimetics or molecular chaperones, to mitigate aggregation risks while preserving translational benefits.

## Acknowledgements

We would like to thank Editage (www.editage.jp) for English language editing. This work was supported by [K.Y.: JSPS KAKENHI Grant Numbers 23K05147 and 19K16259, and S-i.T.: JSPS KAKENHI Grant Number 23K27118].

## Author Contributions

M.M. and H.K. performed the experiments and analyzed the data. K.Y. and S-i.T. supervised the project and interpreted the results. K.Y. wrote the manuscript. K.Y. and S-i.T. reviewed the manuscript. All the authors approved the final version of the manuscript.

## Data Availability

The data supporting the findings of this study are available from the corresponding author upon reasonable request.

---

**Supplementary Figure 1.**
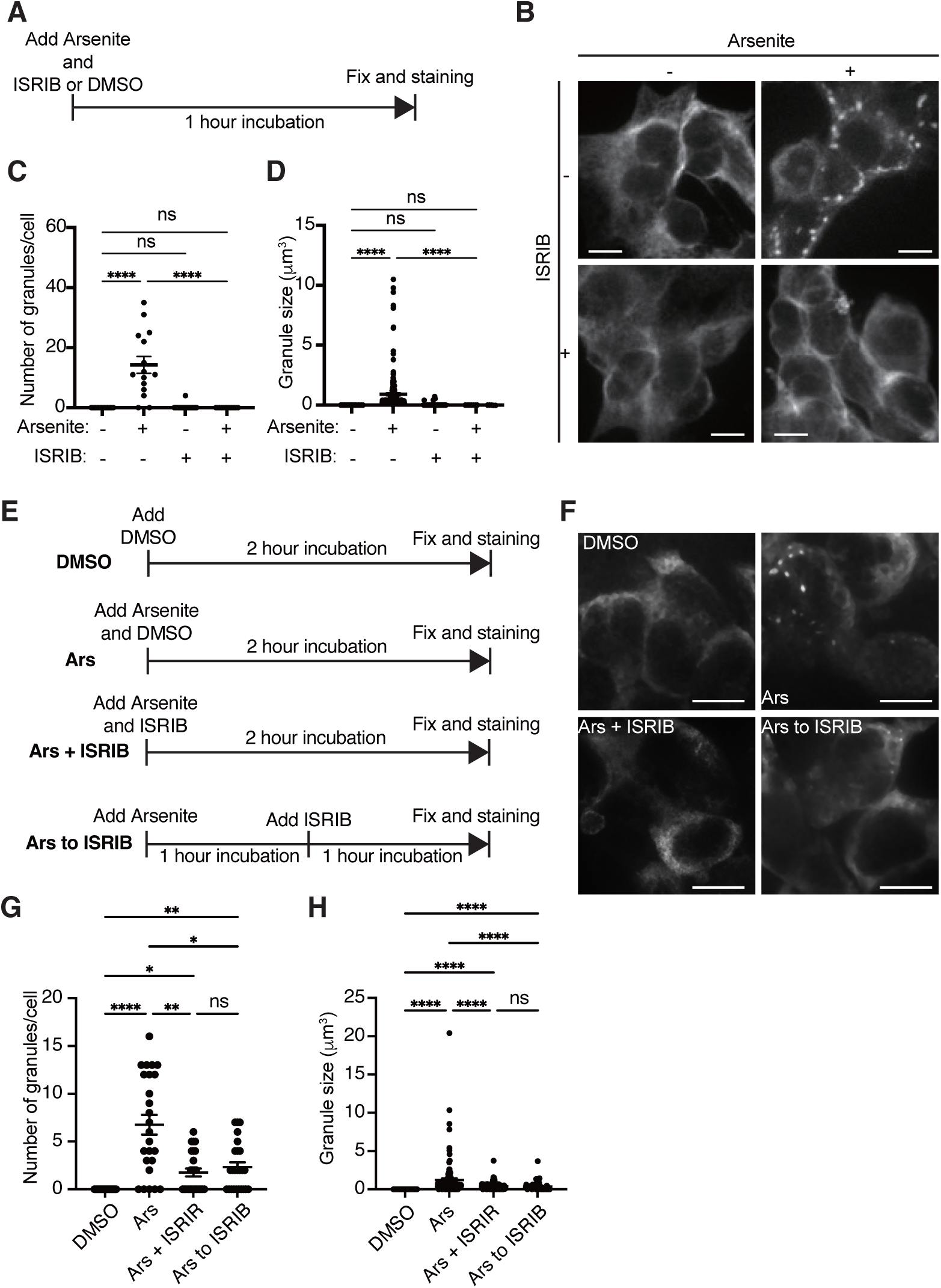
ISRIB dissolves SGs in cells. (**A**) Schematic illustration of ISRIB treatment timing. Cells were treated with sodium arsenite and ISRIB or DMSO simultaneously and incubated for 1 h. (**B**) Representative images of Arsenite- and/or ISRIB-treated HEK293T cells stained with anti-G3BP1 antibody. Bars indicate 10 μm. (**C**) The number of granules in a cell under indicated conditions was counted and plotted. *n(cells)* = 16 (DMSO), 15 (Arsenite+, ISRIB-), 41 (Arsenite-, ISRIB+), and 12 (Arsenite+, ISRIB+) in total. (**D**) The size of granules in a cell in the indicated conditions was counted and plotted. *n* = 16 (DMSO), 215 (Arsenite+, ISRIB-), 44 (Arsenite-, ISRIB+), and 12 (Arsenite+, ISRIB+) in total. Data are from a single trial. Results were compared by using Dunnett’s multiple comparisons test. \*\*\*\**P* < 0.0001. (**E**) Scheme of experimental procedure of ISRIB treatment at different timings. In brief, cells were treated with DMSO (DMSO), arsenite (Ars), or arsenite and ISRIB (Ars + ISRIB) at the same timing and incubated for 2 h. In the case of “Ars to ISRIB”, cells were treated with arsinite for 1 h to form SGs and added ISRIB and further incubated for 1 h. (**F**) Representative images of HEK293T cells treated as labeled. Cells were stained with anti-G3BP1 antibody. Bars indicate 10 μm. (**G**) The number of granules in a cell under indicated conditions was counted and plotted. *n(cells)* = 30 (DMSO), 30 (Ars), 30 (Ars + ISRIB), and 30 (Ars to ISRIB) in total. (**H**) The size of granules in a cell in the indicated conditions was counted and plotted. *n* = 30 (DMSO), 124 (Ars), 89 (Ars + ISRIB), and 73 (Ars to ISRIB) in total. Data are from a single trial (A-D) and three independent trials (E-H). Results were compared by using Dunnett’s multiple comparisons test. \**P* < 0.05, \*\**P* < 0.01 and \*\*\*\**P* < 0.0001.

**Supplementary Figure 2.**
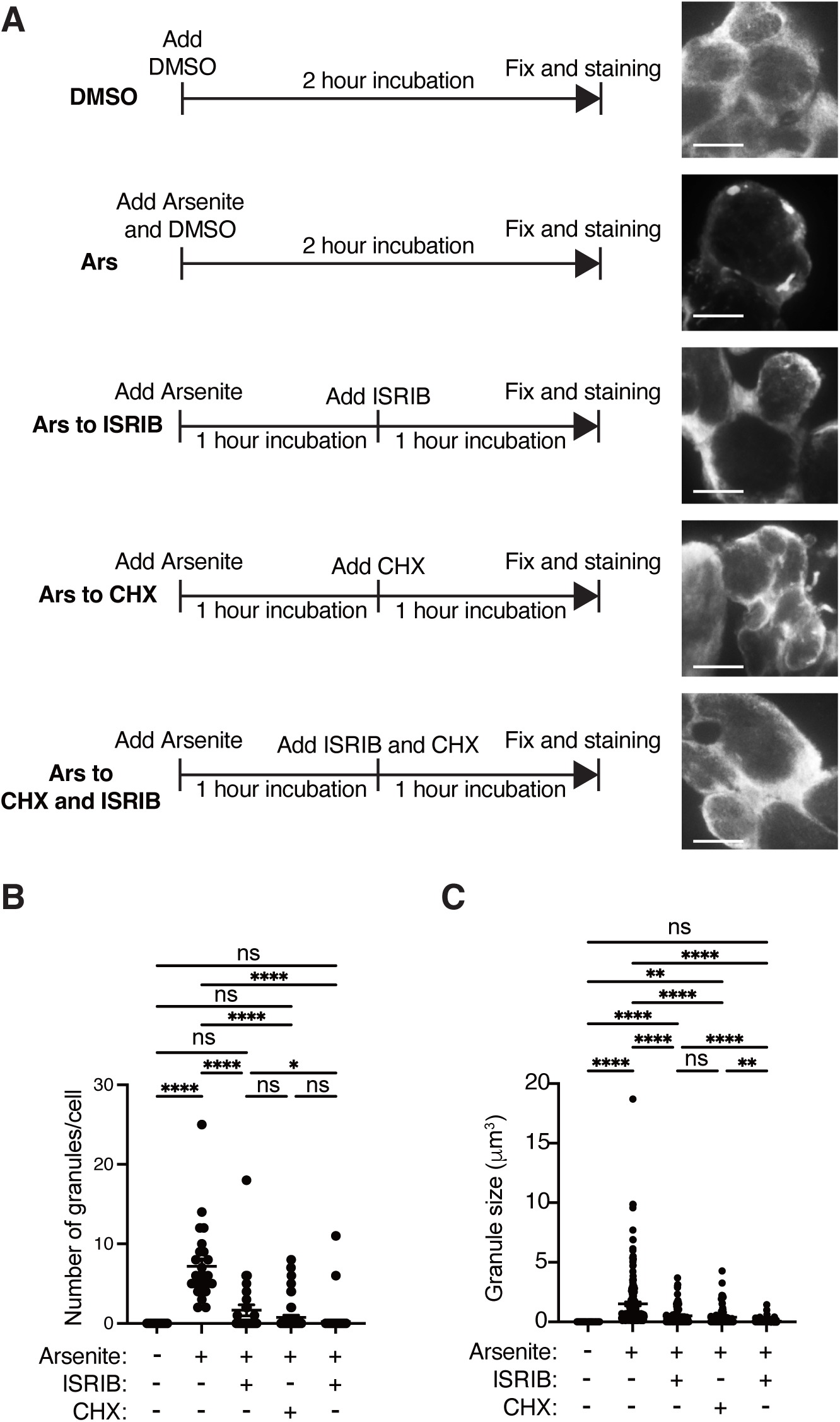
Translational inhibition dissolved SGs, and the effect is enhanced by using ISRIB simultaneously. (**A**) Experimental scheme. HEK293T cells were treated with arsenite for 1 h to induce SGs, followed by the addition of CHX, ISRIB, or both, and incubated for an additional 1 h. Images beside the schemes are the Representative images of cells, which were stained with anti-G3BP1 antibody. Bars indicate 10 μm. (**B**) The number of granules in a cell under indicated conditions was counted and plotted. *n(cells)* = 30 (DMSO), 26 (Ars), 29 (Ars to ISRIB), 50 (Ars to CHX and 79 (Ars to ISRIB and CHX) in total. (**C**) The size of granules in a cell in the indicated conditions was counted and plotted. *n* = 30 (DMSO), 26 (Ars), 29 (Ars to ISRIB), 50 (Ars to CHX and 79 (Ars to ISRIB and CHX) in total. Data are from three independent trials. Results were compared by using Dunnett’s multiple comparisons test. \**P* < 0.05, \*\**P* < 0.01 and \*\*\*\**P* < 0.0001.

**Supplementary Figure 3.**
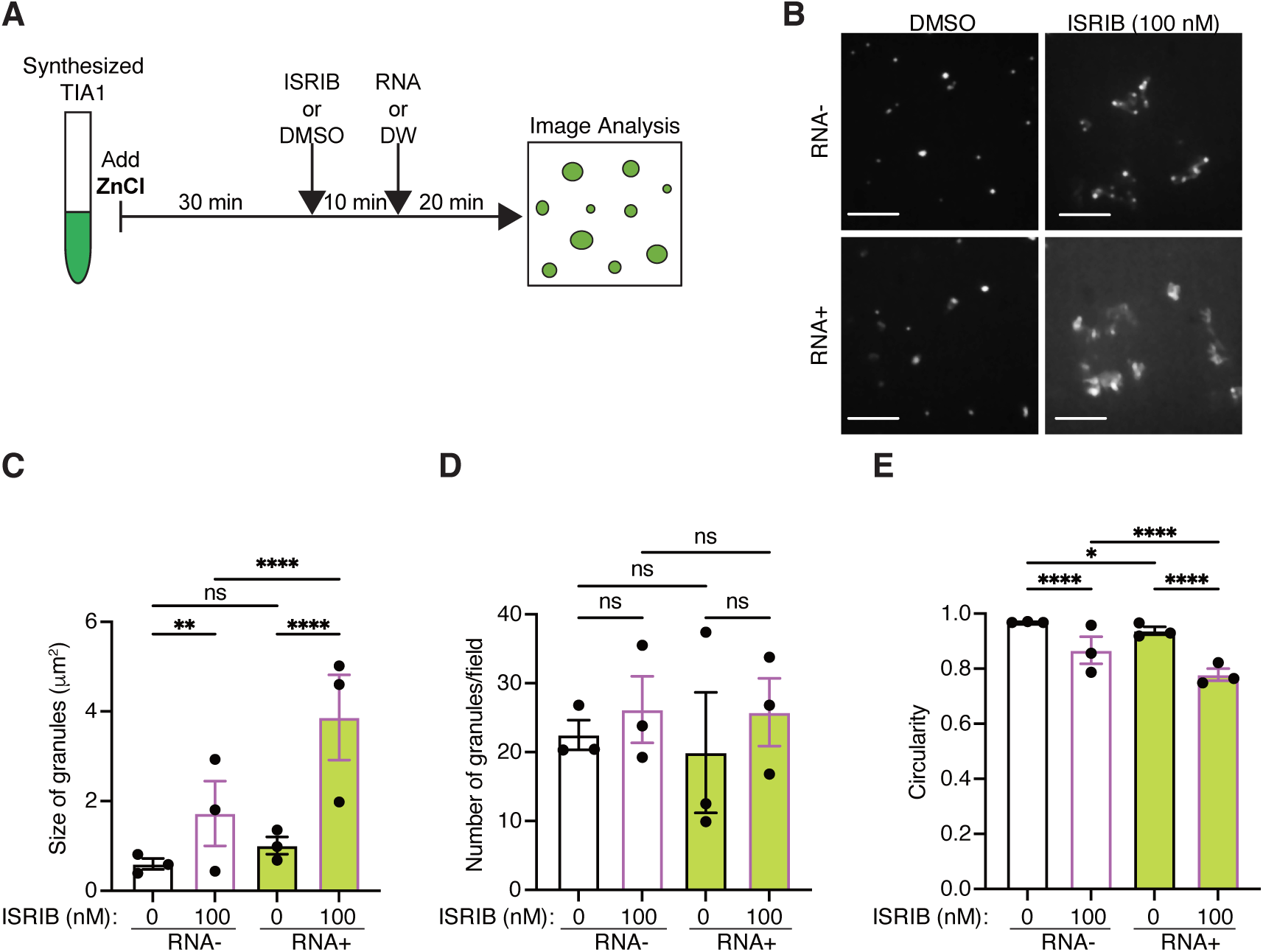
Zn^2+^-induced phase separation of TIA-1 is also affected by ISRIB and is not recovered by RNA addition. **(A)** Schematic view of the experiment. *E. coli-synthesized* and purified mNG-TIA-1 was phase-separated with 3 μM ZnCl^2^. ISRIB was added 30 minutes later, and RNAs were added 10 minutes later. Then samples were observed under a confocal microscope after an additional 20 minutes. **(B)** Representative images of observed mNG-TIA-1 samples treated as in (A). Control samples were treated with DMSO. Bars indicate 10 μm. **(C)** The size of granules analyzed from images of synthesized mNG-TIA-1, treated with (green body) or without RNA (white body), was plotted as a bar graph. Each data point represents the average in one trial. *n(granule)* = 1475 (RNA-, DMSO), 747 (RNA-, 100 nM ISRIB), 548 (RNA+, DMSO), and 774 (RNA+, 100 nM ISRIB) in total. **(D)** The number of granules analyzed from images of synthesized mNG-TIA-1, treated with (green body) or without RNA (white body), was plotted as a bar graph. Each data point represents the average in one trial. *n(granule)* = 30 (RNA-, DMSO), 28 (RNA-, 100 nM ISRIB), 30 (RNA+, DMSO), and 30 (RNA+, 100 nM ISRIB) in total. **(E)** The circularity of granules analyzed from images of synthesized mNG-TIA-1, treated with (green body) or without RNA (white body), was plotted as a bar graph. Each data point represents the average in one trial. *n(granule)* = 1475 (RNA-, DMSO), 747 (RNA-, 100 nM ISRIB), 548 (RNA+, DMSO), and 774 (RNA+, 100 nM ISRIB) in total. All data points were collected from independent trials. Results were compared by using Dunnett’s multiple comparisons test. \*\*\*\**P* < 0.0001.

